# Dissection of purified LINE-1 reveals distinct nuclear and cytoplasmic intermediates

**DOI:** 10.1101/157818

**Authors:** K.R. Molloy, M.S. Taylor, I. Altukhov, P. Mita, H. Jiang, E.M. Adney, A. Wudzinska, D. Ischenko, K.H. Burns, D. Fenyö, B.T. Chait, D. Alexeev, M.P. Rout, J.D. Boeke, J. LaCava

## Abstract

Long Interspersed Nuclear Element-1 (LINE-1, L1) is a mobile genetic element active in human genomes. L1-encoded ORF1 and ORF2 proteins bind L1 RNAs, forming ribonucleoproteins (RNPs). These RNPs interact with diverse host proteins, some repressive and others required for the L1 lifecycle. Using differential affinity purifications and quantitative mass spectrometry, we have characterized the proteins associated with distinctive L1 macromolecular complexes. Our findings support the presence of multiple L1-derived retrotransposition intermediates *in vivo*. Among them, we describe a cytoplasmic intermediate that we hypothesize to be the canonical ORF1p/ORF2p/L1*-*RNA-containing RNP, and we describe a nuclear population containing ORF2p, but lacking ORF1p, which likely contains host factors participating in template-primed reverse transcription.

## 3. Introduction

Sequences resulting from retrotransposition constitute more than half of the human genome and are considered to be major change agents in eukaryotic genome evolution (Kazazian, 2004). L1 retrotransposons have been particularly active in mammals (Furano *et al*, 2004), comprising ~20% of the human genome (Lander *et al*, 2001); somatic retrotransposition has been widely implicated in cancer progression (Lee *et al*, 2012; Tubio *et al*, 2014) and may even play a role in neural development (Muotri *et al*, 2005). Despite the magnitude of their contributions to mammalian genomes, L1 genes are modest in size. A full-length L1 transcript is ~6 knt long and functions as a bicistronic mRNA that encodes two polypeptides, ORF1p and ORF2p (Ostertag & Kazazian, 2001), which respectively comprise a homotrimeric RNA binding protein with nucleic acid chaperone activity (Martin & Bushman, 2001) and a multifunctional protein with endonuclease and reverse transcriptase activities (Mathias *et al*, 1991; Feng *et al*, 1996). Recently, a primate-specific third ORF, named *ORF0*, has been identified on the Crick strand of the L1 gene; this ORF encodes a 71 amino acid peptide and may generate insertion site-dependent ORFs via splicing (Denli *et al*, 2015). ORF1p and ORF2p are thought to interact preferentially with the L1 RNA from which they were translated (in cis), forming a ribonucleoprotein (RNP) (Kulpa & Moran, 2006; Taylor *et al*, 2013) considered to be the canonical direct intermediate of retrotransposition (Hohjoh & Singer, 1996; Kulpa & Moran, 2005; Martin, 1991; Kulpa & Moran, 2006; Doucet *et al*, 2010). L1s also require host factors to complete their lifecycle (Suzuki *et al*, 2009; Peddigari *et al*, 2013; Dai *et al*, 2012; Taylor *et al*, 2013) and, consistent with a fundamentally parasitic relationship (Beauregard *et al*, 2008), the host has responded by evolving mechanisms that suppress retrotransposition (Goodier *et al*, 2013; Arjan-Odedra *et al*, 2012; Goodier *et al*, 2012; Niewiadomska *et al*, 2007). It follows that as the host and the parasite compete, L1 expression is likely to produce a multiplicity of RNP forms engaged in discrete stages of retrotransposition, suppression, or degradation.

Although L1 DNA sequences are modestly sized compared to typical human genes, L1 intermediates are nevertheless RNPs with a substantially sized RNA component; e.g. larger than the ~5 knt 28S rRNA (Gonzalez *et al*, 1985) and approximately three to four times the size of a "typical" mRNA transcript (Lander *et al*, 2001; Sommer & Cohen, 1980). Therefore, it is likely that many proteins within L1 RNPs form interactions influenced directly and indirectly by physical contacts with the L1 RNA. We previously determined that L1 RNA comprised ~20-30% of mappable RNA-seq reads in ORF2p-3xFLAG affinity captured fractions (Taylor *et al*, 2013). We also observed that the retention of ORF1p and UPF1 within affinity captured L1 RNPs was reduced by treatment with RNases (Taylor *et al*, 2013). In the same study we observed that two populations of ORF2p-associated proteins could be separated by split-tandem affinity capture (ORF2p followed by ORF1p), a two-dimensional affinity enrichment procedure (Caspary *et al*, 1999; Taylor *et al*, 2013). Initial characterization of these two L1 populations by western blotting suggested that discrete L1 populations were likely primed for function in different stages of the lifecycle. We therefore expected additional uncharacterized complexity in the spectrum of L1-associated complexes present in our affinity purified fractions.

In this study, we have used quantitative mass spectrometry (MS) to investigate the proteomic characteristics of endogenously assembled ectopic L1 complexes present in an assortment of affinity-enriched fractions. We revisited RNase treatment and split-tandem affinity capture approaches and complemented them with in-cell localization of ORF proteins by immunofluorescence microscopy (see also the companion manuscript by **Mita et al. [co-submission]**). We additionally explored L1 proteomes **associated with catalytically-** inactivated ORF2p point mutants and monitored the rates of protein exchange from L1s *in vitro.* Taken together, our data support the existence of a variety of putative L1-related protein complexes.

## 4. Results

Affinity proteomic experiments conducted in this study use quantitative MS based upon metabolic labeling (Oda *et al*, 1999). Two main experimental designs (and modifications thereof) facilitating quantitative cross-sample comparisons have been used: SILAC (Ong *et al*, 2002; Wang & Huang, 2008) and I-DIRT (Tackett *et al*, 2005; Taylor *et al*, 2013). In these approaches, cells are grown for several doublings in media containing amino acids composed either of naturally-occurring ’light’ isotopes or biologically identical ’heavy’ isotopes (e.g. ^13^C, ^15^N lysine and arginine), such that the proteomes are thoroughly labeled. Protein fractions derived from the differently labeled cell populations, obtained e.g. before and after experimental manipulations are applied, are mixed and the relative differences in proteins contributed by each fraction are precisely measured by mass spectrometry. In addition to the above cited studies, these approaches have been adapted to numerous biological questions using a variety of analytical frameworks e.g. (Byrum *et al*, 2011; Luo *et al*, 2016; Trinkle-Mulcahy *et al*, 2008; Ohta *et al*, 2010; Kaake *et al*, 2010; Geiger *et al*, 2011). Because it is challenging to speculate on the potential physiological roles of protein interactions that form after extraction from the cell, we often use I-DIRT, which allows the discrimination of protein-protein interactions formed in-cell from those occurring post-extraction. Our prior affinity proteomic study, based on I-DIRT, identified 37 putative *in vivo* interactors (Taylor *et al*, 2013), described in **Table 1**. Therefore, primarily focus here on the behaviors of these !I-DIRT significant" L1 interactors, in order to determine their molecular associations and ascertain the variety of distinctive macromolecular complexes formed in-cell that co-purify with affinity-tagged ORF2p. The complete lists of proteins detected in each experiment are presented in the supplementary information (see **Table S1**). Except where noted otherwise, the presented experiments were conducted in suspension-cultured HEK-293T_LD_ cells, using a synthetic L1 construct - *ORFeus*-HS – driving the expression 3xFLAG-tagged L1 (*ORF1*; *ORF2::3xFLAG*; 3’-UTR) from a tetracycline inducible minimal-CMV promoter, harbored on a mammalian episome (pLD401 (Taylor *et al*, 2013; An *et al*, 2011; Dai *et al*, 2012)). All purified L1s described in this study were obtained by affinity capture of ORF2p-3xFLAG before further experimental manipulations were applied. We have represented any ambiguous protein group, which occurs when the same peptides identify a group of homologous protein sequences, with a single, consistently applied gene symbol and a superscript ’a’ in all figures. **Table S1** contains the references to other proteins explaining the presence of the same peptides. For example, RPS27A, (ubiquitin) UBB, UBC, and (ribosomal Protein L40) UBA52 can be explained by common ubiquitin peptides shared by these genes. RPS27A-specific peptides were not identified in this study, but we retained the nomenclature for consistency with our previous work; HSPA1A is reported in this study, but cannot be distinguished from the essentially identical protein product of HSPA1B.

**Table 1:** *Putative L1 interactors*: Through a series of affinity capture experiments using I-DIRT, we characterized a set of putative host-encoded L1 interactors (Taylor *et al*, 2013). The proteins observed were associated with both ORF1p and ORF2p (highlighted in blue), or only with only one ORF protein. Proteins only observed in association with ORF2p are highlighted in magenta. The two highlighted populations are the central focus of this study.

| **Gene Symbol** | **Uniprot Symbol** | **Protein** | **co-IP** |
| --- | --- | --- | --- |
| L1RE1 | Q9UN81 | ORF1p | ORF1/2 |
| N/A | O00370 | ORF2p | ORF1/2 |
| MOV10 | Q9HCE1 | Putative helicase MOV-10 | ORF1/2 |
| PABPC1 | P11940 | Polyadenylate-binding protein 1 | ORF1/2 |
| PABPC4 | Q13310 | Polyadenylate-binding protein 4 | ORF1/2 |
| UPF1 | Q92900 | Regulator of nonsense transcripts 1 | ORF1/2 |
| ZCCHC3 | Q9NUD5 | Zinc finger CCHC domain-containing protein 3 | ORF1/2 |
| FKBP4 | Q02790 | Peptidyl-prolyl cis-trans isomerase FKBP4 | ORF2 |
| HAX1 | O00165 | HCLS1-associated protein X-1 | ORF2 |
| HMCES | Q96FZ2 | Embryonic stem cell-specific 5-hydroxymethylcytosine-binding protein | ORF2 |
| HSP90AA1 | P07900 | Heat shock protein HSP 90-alpha | ORF2 |
| HSP90AB1 | P08238 | Heat shock protein HSP 90-beta | ORF2 |
| HSPA1A | P0DMV8 | Heat shock 70 kDa protein 1A | ORF2 |
| HSPA8 | P11142 | Heat shock cognate 71 kDa protein | ORF2 |
| IPO7 | O95373 | Importin-7 | ORF2 |
| NAP1L1 | P55209 | Nucleosome assembly protein 1-like 1 | ORF2 |
| NAP1L4 | Q99733 | Nucleosome assembly protein 1-like 4 | ORF2 |
| PARP1 | P09874 | Poly [ADP-ribose] polymerase 1 | ORF2 |
| PCNA | P12004 | Proliferating cell nuclear antigen | ORF2 |
| PURA | Q00577 | Transcriptional activator protein Pur-alpha | ORF2 |
| PURB | Q96QR8 | Transcriptional activator protein Pur-beta | ORF2 |
| RPS27A | P62979 | Ubiquitin-40S ribosomal protein S27a | ORF2 |
| TIMM13 | Q9Y5L4 | Mitochondrial import inner membrane translocase subunit Tim13 | ORF2 |
| TOP1 | P11387 | DNA topoisomerase 1 | ORF2 |
| TOMM40 | O96008 | Mitochondrial import receptor subunit TOM40 homolog | ORF2 |
| TUBB | P07437 | Tubulin beta chain | ORF2 |
| TUBB4B | P68371 | Tubulin beta-4B chain | ORF2 |
| YME1L | Q96TA2 | ATP-dependent zinc metalloprotease YME1L1 | ORF2 |
| CORO1B | Q9BR76 | Coronin-1B | ORF1 |
| DDX6 | P26196 | Probable ATP-dependent RNA helicase DDX6 | ORF1 |
| ERAL1 | O75616 | GTPase Era, mitochondrial | ORF1 |
| HIST1H2BO | P23527 | Histone H2B type 1-O | ORF1 |
| LARP7 | Q4G0J3 | La-related protein 7 | ORF1 |
| MEPCE | Q7L2J0 | 7SK snRNA methylphosphate capping enzyme | ORF1 |
| PABPC4L | P0CB38 | Polyadenylate-binding protein 4-like | ORF1 |
| TROVE2 | P10155 | 60 kDa SS-A/Ro ribonucleoprotein | ORF1 |
| YARS2 | Q9Y2Z4 | Tyrosine--tRNA ligase, mitochondrial | ORF1 |

### 4.1 RNase-sensitivity exhibited by components of affinity purified L1 RNPs

**Figure 1 (panels A-C)** illustrates the approach and displays the findings of our assay designed to reveal which proteins depend upon the presence of intact L1 RNA for retention within the purified L1 RNPs. Briefly, metabolically-labeled affinity captured L1s were treated either with a mixture of RNases A and T1 — thus releasing proteins that require intact RNA to remain linked to ORF2p and the affinity medium — or BSA, as an inert control. After removing the fractions released by the RNase or BSA treatments, the proteins remaining on the affinity media were eluted with lithium dodecyl sulfate (LDS), mixed together, and then analyzed by MS. Proteins released, and so depleted, by RNase treatment were thus found to be more abundant in the BSA-treated control. The results obtained corroborate and extend our previous findings: ORF1p and UPF1 exhibited RNase-sensitivity (Taylor *et al*, 2013). We also observed that ZCCHC3 and MOV10 exhibited RNase-sensitivity to a level similar to ORF1p. The remaining I-DIRT significant proteins were RNase-resistant in this assay. With the exception of the PABPC1/4 proteins (and ORF2p itself, **see Discussion**), the I-DIRT significant proteins (**colored nodes, Fig. 1C**) that were resistant to RNase treatment (nearest the origin of the graph) classify ontologically as nuclear proteins (GO:0005634, p ≈ 3 × 10^−4^, **see Methods**). These same proteins were previously observed as specific L1 interactors in I-DIRT experiments targeting ORF2p but not in those targeting ORF1p; in contrast, the proteins that demonstrated RNase-sensitivity: ORF1, MOV10, ZCCHC3, and UPF1 were observed in both ORF1p and ORF2p I-DIRT experiments (**Table 1**). Stated another way, the proteins released upon treating an affinity captured ORF2p fraction with RNases are among those that can also be obtained when affinity capturing ORF1p directly, while those that are RNase-resistant are not ORF1p interactors (Taylor *et al*, 2013). The ORF1p-linked, I-DIRT significant, RNase-sensitive proteins were too few to obtain a high confidence assessment of ontological enrichment; but, when combined with remaining proteins exhibiting sensitivity to the RNase treatment (**black nodes, Fig. 1C**), they together classified as ’RNA binding’ (GO:0003723, p ≈ 1 × 10^−11^). This analysis also revealed a statistically significant overrepresentation of genes associated with the exon junction complex (EJC, GO: 0035145, p ≈ 1 × 10^−6^, discussed below). ORF1p/2p-associated interactomes appear to overlap on the L1 RNA, as expected. However, considering the nuclear ontology associated with the RNase-resistant ORF2p interactors, we conclude that ORF1p probably does not interact with ORF2p in the nucleus, where the ORF2p interactome is either not broadly dependent upon L1 RNA, or is protected from RNases. ORF1p may be displaced from the L1 RNA during the nuclear portion of the lifecycle. Thus, host-encoded proteins appear to segregate themselves into groups according to their L1 ORF protein associations, and that these groups respond differently, yet in an apparently coordinated fashion, to RNase treatment. This observation led to the hypothesis that our ORF2p-3xFLAG affinity purified L1s constitute a composite purification of at least, but not limited to, (1) a cytoplasmic population of L1-RNA-dependent, ORF1p/ORF2p-containing L1 RNPs, and (2) an ORF1p-independent nuclear population associated with ORF2p.

**Figure 1.**
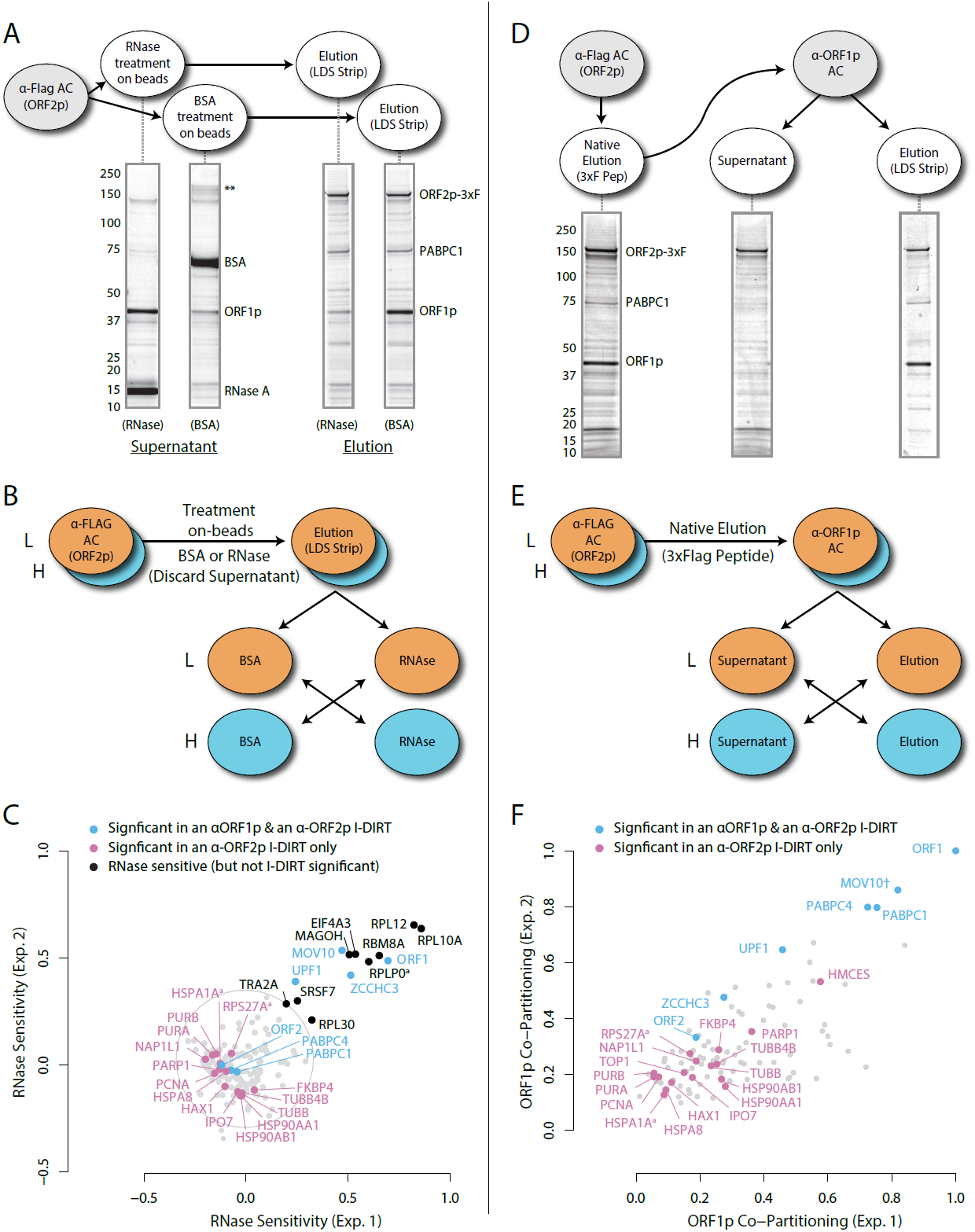
(**A)** *On-bead RNase-sensitivity assay*: L1 complexes were affinity captured by ORF2p-3xFLAG. The magnetic media were then treated with a solution containing either a mixture of RNases A and T1 or BSA. After treatment the supernatants were removed and the remaining bound material was released by treatment with LDS. Proteins requiring intact RNA to maintain stable interactions with immobilized ORF2p were released from the RNase-treated medium, while the BSA-treated sample provided a control for the spontaneous release of proteins from the medium over the time of the assay. Representative SDS-PAGE / Coomassie blue stained gel lanes are shown for each fraction. (**B**) The experiments described above was carried out in duplicate, once with light isotopically labeled cells (L) and once with heavy isotopically labeled cells (H), resulting in four label-swapped, SILAC duplicates (one light set & one heavy set). The four fractions were cross-mixed and the differential protein retention upon the affinity medium during the treatments (BSA vs. RNase) was assessed by quantitative MS. (**C**) Results from the RNase-sensitivity assay graphed as the fraction of each detected protein present in the BSA-treated sample (RNase-sensitive proteins are *more* present in the BSA treated sample), normalized such that proteins that did not change upon treatment with RNases are centered at the origin. A cut-off of p = 10^−3^ for RNase-sensitivity is indicated by a light gray circle; proteins that are RNase-sensitive with a statistical significance of p < 10^−3^ are outside the circle. Proteins previously ranked significant by I-DIRT analysis (**Table 1**) are labeled and displayed in blue or magenta (as indicated); black nodes were not found to be significant by I-DIRT but were labeled if found to be RNase-sensitive; gray, unlabeled nodes were not found to be significant by I-DIRT. (**D**) *Split-tandem affinity capture*: L1 complexes were affinity captured by ORF2p-3xFLAG. After native elution with 3xFLAG peptide, this fraction was subsequently depleted of ORF1p containing complexes using an α-ORF1 conjugated magnetic medium, resulting in a supernatant fraction depleted of ORF1p-containing complexes. The α-ORF1 bound material was then released with LDS, yielding an elution fraction enriched for ORF1p-containing complexes. Representative SDS-PAGE / Coomassie blue stained results for each fraction are shown. (**E**) SILAC duplicates, two supernatants and two elution, were cross-mixed to enable an assessment of the relative protein content of each fraction by quantitative MS. (**F**) The results from split-tandem affinity capture graphed as the fraction of each protein observed in the elution sample. In order to easily visualize the relative degree of co-partitioning of constituent proteins with ORF1p, these data were normalized, setting the fraction of ORF1p in the elution to 1. Proteins which were previously ranked significant by I-DIRT analysis are labeled and displayed in blue or magenta (as indicated); gray, unlabeled nodes were not found to be significant by I-DIRT. MOV10 is marked with a dagger because in one replicate of this experiment it was detected by a single unique peptide, whereas we have enforced a minimum of two peptides (see **Methods**) for all other proteins, throughout all other proteomic analyses presented here.

While effects of PABPC1, MOV10, and UPF1 on L1 activity have been described (Arjan-Odedra *et al*, 2012; Taylor *et al*, 2013; Dai *et al*, 2012), effects of ZCCHC3 on L1 remained uncharacterized. ZCCHC3 is an RNA-binding protein associated with poly(A)+ RNAs (Castello *et al*, 2012) but otherwise little is known concerning its functions. Notably, in a genome-wide screen, small interfering (si)RNA knockdown of ZCCHC3 was observed to increase the infectivity of the Hepatitis C, a positive sense RNA virus (Li *et al*, 2009); and ZCCHC3 was observed to copurify with affinity captured HIV, a retrovirus, at a very high SILAC ratio (>10), supporting the specificity of this interaction (Engeland *et al*, 2014). We therefore explored the effects on L1 mobility both of over-expression and siRNA knockdown of ZCCHC3. Over-expression of ZCCHC3 reduced L1 retrotransposition to ~10% that observed in the control, consistent with a negative regulatory role for ZCCHC3 in the L1 lifecycle; small interfering RNA (siRNA) knockdown of ZCCHC3 induced a modest increase in retrotransposition compared to a scrambled control siRNA (~1.9× ±0.1; **Table S2**). Moreover, although not among our I-DIRT hits (see **Discussion**), the presence of EJC components (MAGOH, RBM8A, EIF4A3, UPF1) among the RNase-sensitive fraction of proteins intrigued us, given that L1 genes are intronless. We speculated that L1s may use EJCs to enhance nuclear export, evade degradation by host defenses, and/or aggregate with mRNPs within cytoplasmic granules. For this reason we carried out a series of siRNA knockdowns of these EJC components and other physically or functionally related proteins found in the affinity purified fraction (listed in **Table S2**). siRNA knockdowns of RBM8A and EIF4A3 caused inviability of the cell line. We found that knocking-down MAGOH and the EJC-linked protein IGF2BP1 (Jønson *et al*, 2007) both, respectively, reduced retrotransposition by ~50%, consistent with a role in L1 proliferation; although, these knockdowns also caused a reduction in viability of the cell line (**See Discussion**).

### 4.2 Split-tandem separation of compartment-specific L1 ORF-associated complexes

To further test our hypothesis and better characterize the components of our L1 fraction, we conducted split-tandem affinity capture. **Figure 1 (panels E-F)** illustrates the approach and displays the findings of the assay, which physically separated ORF1p/ORF2p-containing L1 RNPs from a presumptive ’only-ORF2p-associated’ population. Briefly, metabolically-labeled L1s were affinity captured by ORF2p-3xFLAG (1^st^ dimension) and the obtained composite was subsequently further fractionated by α-ORF1p affinity capture (2^nd^ dimension, or split-tandem purification), resulting in α-ORF1p-bound and unbound fractions. The bound fraction was eluted from the affinity medium with LDS. The bound and unbound fractions were then mixed and analyzed by MS to ascertain proteomic differences between them. The fraction eluted from the α-ORF1p medium contained the population of proteins physically linked to *both* ORF2p and ORF1p, whereas the supernatant from the α-ORF1p affinity capture contained the proteins associated *only* with ORF2p (and, formally, those which have dissociated from the ORF1p/ORF2p RNP). The results corroborated our previous observations that: i) almost all of the ORF1p partitioned into the elution fractions, ii) a quarter of the ORF2p (~26%) followed ORF1p during the α-ORF1p affinity capture, iii) roughly half of the UPF1 (~55%) followed ORF1p, and iv) most of the PCNA (~87%) remained in the ORF1p-depleted supernatant fraction (**Fig. 1F**, and consistent with prior estimates based on protein staining and western blotting (Taylor *et al*, 2013)); thus v) supporting the existence of at least two distinct populations of L1-ORF-protein-containing complexes in our affinity purifications.

The population eluted from the α-ORF1p affinity medium (**Fig. 1D,** far right gel lane,and nodes located in the upper right of the graph, **panel F)** is consistent with the composition of the ORF1p/ORF2p-containing L1 RNP suggested above. Our split-tandem separation segregated the constituents of the purified L1 fraction comparably to the RNase-sensitivity assay, both in terms of which proteins co-segegate with ORF1p/ORF2p (compare **Fig. 1C** and **F,** blue nodes, upper right of graphs) as well as those which appear to be linked only to ORF2p (compare Figs. **1C** and **F**, magenta nodes, lower left of the graphs). The ORF1p/ORF2p RNPs obtained by split-tandem purification include putative *in vivo* interactions associated with both - α ORF1p and α-ORF2p I-DIRT affinity capture experiments; whereas the unbound, ORF1p-independent fraction includes proteins previously observed as significant only in α-ORF2p I-DIRT experiments (**Table 1**). Analysis of the nodes whose degree of ORF1p association was similar to that of UPF1 (**blue nodes exhibiting ≥ 55% ORF1p co-partitioning**, **Fig. 1F**) revealed that they map ontologically to a ’cytoplasmic ribonucleoprotein granule’ classification (GO:0036464, p ≈ 6 × 10^−8^; **see Discussion**). In contrast, all sixteen proteins exhibiting ORF1p co-partitioning approximately equal to or less than that of ORF2p were predominantly found in the supernatant fraction and were enriched for cell-compartment-specific association with the nucleus (GO:0005634, p ≈ 4 × 10^−5^ **Fig. 1F, all magenta nodes below PARP1, inclusive, ≤ 36%**). These two fractions therefore appear to be associated with different cell compartments, reaffirming our postulate: the ORF1p/ORF2p-containing population is a cytoplasmic intermediate related to the canonical L1 RNP typically ascribed to L1 assembly in the literature, and the predominantly ORF2p-associated population comprises a putative nuclear interactome; each therefore referred to, respectively, as cytoplasmic and nuclear L1 interactomes hereafter.

From the same analysis, we noted that PURA, PURB, PCNA, and TOP1 which all partition predominantly with nuclear L1, exhibited an ontological co-enrichment (termed ’nuclear replication fork,’ GO:0043596, p ≈ 3 × 10^−4^). The nodes representative of PURA, PURB, and PCNA appeared to exhibit a striking proximity to one another, suggesting highly similar co-fractionation behavior potentially indicative of direct physical interactions. In an effort to examine this possibility, we graphed the frequency distribution of the proximities of all three-node-clusters observed within **Figure 1F**, revealing the likelihood of the PURA/PURB/PCNA cluster to be p = 3.2 × 10^−7^ (see **Supplementary Methods**). We therefore concluded that PURA, PURB, PCNA, and (perhaps at a lower affinity) TOP1, likely constitute a physically associated functional module interacting with L1. In further support of this assertion, we noted that known functionally linked protein pairs PABPC1/PABPC4 (cytoplasmic) (Jønson *et al*, 2007; Katzenellenbogen *et al*, 2007) and HSPA8/HSPA1A (nuclear) (Jønson *et al*, 2007; Nellist *et al*, 2005) also exhibited comparable co-partitioning by visual inspection, and statistical testing of these clusters revealed the similarity of their co-partitioning to be significant at p ≈ 0.001 for the former, and p ≈ 0.0002 for the latter. The observed variation in co-partitioning behavior between the different proteins comprising the nuclear L1 fraction might reflect the presence of multiple distinctive (sub)complexes present within this population.

To validate our hypothesis that these proteins are associated with ORF2p in the nucleus, we carried out ORF2p-3xFLAG affinity capture from chromatin-enriched sub-cellular fractions and found that the co-purified proteins we identified largely overlapped with those described above (**Table S3**), including: PARP1, PCNA, UPF1, PURA, and TOP1. Although we observed that silencing PCNA expression adversely affects L1 retrotransposition (Taylor *et al*, 2013), we found that knocking down TOP1 approximately doubled retrotransposition frequency, while a more modest 1.4x increase effect was observed for PURA, and no substantial effect was observed for PURB, compared to a scrambled siRNA control. In contrast, over-expression of PURA reduced retrotransposition to ~20% of the expected level (**Table S2**).

### 4.3 ORF1p/ORF2p immunofluorescence protein localization

Although our proteomic and biochemical analyses supported the existence of distinctive nuclear and cytoplasmic L1 populations, our prior immunofluorescence (IF) analyses did not reveal an apparent nuclear population, leading us to revisit IF studies. Previously, IF of ORF1p and ORF2p in HeLa and HEK-293T cells yielded two striking observations: i) ORF2 expression was seemingly stochastic, with ORF2p observed in ~30% of cells; and ii) while ORF1p and ORF2p co-localized in cells that exhibited both, we did not observe an apparent nuclear population of either protein (Taylor *et al*, 2013). Subsequently, we noted an absence of mitotic cells from these preparations. Reasoning that these cells were lost due to selective adherence on glass slides, and noting that cell division has been reported to be promote L1 transposition (Xie *et al*, 2013; Shi *et al*, 2007), we repeated the assays using puromycin-selected Tet-on HeLa cells grown on fibronectin coated coverslips. The results are shown in **Figure 2**.

**Figure 2.**
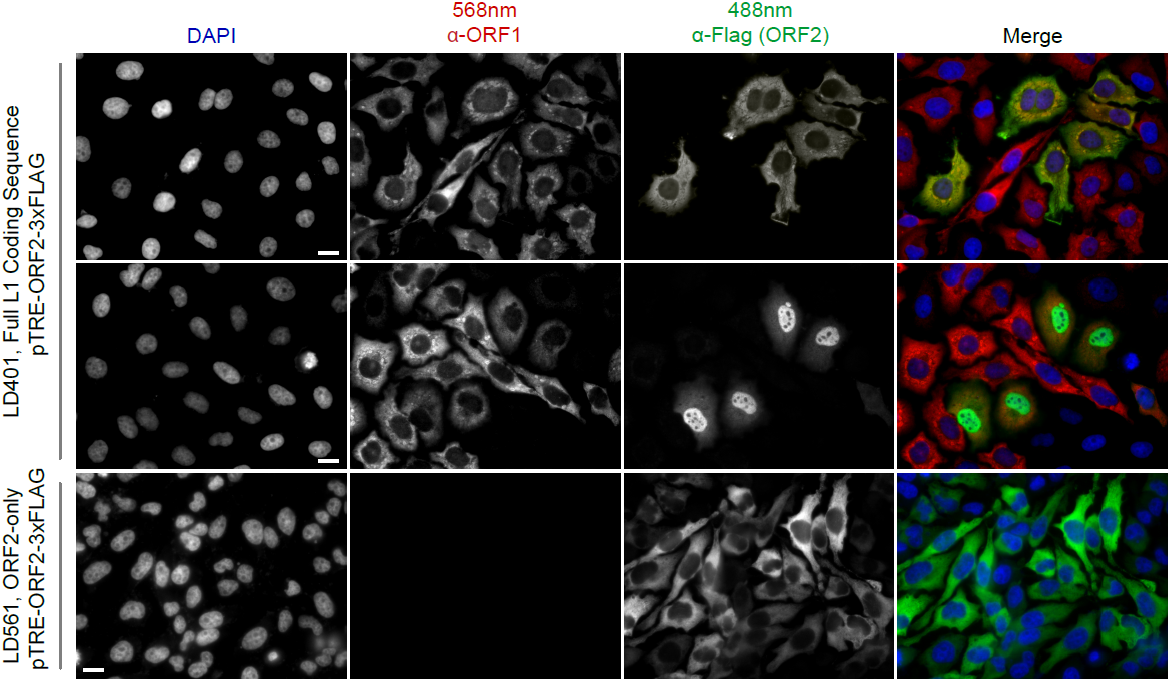
*Immunofluorescent Imaging Reveals ORF1p is Required for Nuclear ORF2p*: Puromycin-selected HeLa-M2 cells containing pLD401 (Tet promoter, [*ORFeus*-Hs] full L1 coding sequence, ORF2p-3xFLAG, top two rows) or pLD561 (Tet promoter, *ΔORF1*, ORF2p-3xFLAG, bottom row) were plated on fibronectin-coated coverslips and induced for 24 hr with doxycycline prior to fixation and staining. With pLD401, the previously-observed pattern of cytoplasmic-only ORFs (top row) and a new pattern of pairs of cells expressing ORF2p in the nucleus (middle row) were apparent. When ORF1p is omitted from the construct (pLD561, bottom row), nuclear ORF2p was not apparent. Scale bars: 10 micrometers.

The modified IF assay corroborated our prior results in that nearly all the cells exhibited cytoplasmic ORF1p and ~30% also exhibited cytoplasmic ORF2p, which co-localized with ORF1p (**Fig. 2, top row**). However, a new population of cells also became apparent, consisting of pairs of cells exhibiting nuclear localized ORF2p (**Fig. 2, middle row**). These cells accounted for <10% of the population and, because of they always occurred in proximal pairs, are presumed to have recently gone through mitosis. Expression of *ORF2* in the absence of *ORF1* (*ΔORF1*; pLD561) resulted in >90% of cells exhibiting cytoplasmic ORF2p, consistent with our previous work (Taylor *et al*, 2013). We did not observe instances of nuclear ORF2p using the construct (**Fig. 2, bottom row**), suggesting that ORF1p is required for ORF2p nuclear localization. In a separate study, including more detailed analyses ORF protein localization, **Mita et al. (co-submission)** observed that both ORF proteins enter the nucleus of HeLa cells during mitosis, however, nuclear ORF1p does not seem to be physically associated with nuclear ORF2p (**see Discussion**). Taken together, the data obtained from the modified IF experiments aligned well with our proteomic and biochemical data: L1 expression resulted in *at least* two distinct populations: cytoplasmic complexes containing both ORF1p and ORF2p, and nuclear complexes containing ORF2p while potentially lacking ORF1p.

### 4.4 The effects of retrotransposition-blocking point mutations on the interactomes of purified L1 RNPs

Based on the hypothesis that our composite purifications contain bona fide nuclear intermediates, we decided to explore the effects of catalytic point mutations within the ORF2p endonuclease and reverse transcriptase domains, respectively. We reasoned that such mutants may bottleneck L1 intermediates at the catalytic steps associated with host gDNA cleavage and L1 cDNA synthesis, potentially revealing protein associations that are important for these discrete aspects of template primed reverse transcription (TPRT), the presumed mechanism of L1 transposition (Luan *et al*, 1993; Feng *et al*, 1996; Cost *et al*, 2002). For this we used an H230A mutation to inactivate the endonuclease activity (EN^-^ / pLD567), and a D702Y mutation to inactivate the reverse transcriptase activity (RT^-^ / pLD624) (Taylor *et al*, 2013). **Figure 3** illustrates the approach and displays the findings of our assay. Broadly, our findings revealed several classes of behaviors (**Fig. 3B**). Two classes of behavior appeared to be particularly striking: (1) the yield of constituents of cytoplasmic L1s was reduced, relative to WT, by the EN mutation, yet elevated by the RT mutation (**Fig. 3B, left side**); and (2) numerous constituents of nuclear L1s were elevated in yield by the EN mutation but reduced or nominally unchanged, relative to WT, by the RT mutation (**Fig. 3B, right side**). While reductions in the levels of cytoplasmic L1 proteins within EN^-^ L1 RNPs may be difficult to rationalize, accumulation of these proteins within RT^-^ L1s could be explained by a decrease in the frequency that ORF2p traverses the L1 RNA, normally displacing these proteins to some degree even within the cytoplasm. With respect to the second group, IPO7, NAP1L4, NAP1L1, FKBP4, HSP90AA1, and HSP90AB1 were all elevated in the EN^-^ mutants, potentially implicating these proteins as part of an L1 complex (or complexes) immediately preceding DNA cleavage. Notably, there is a third class of proteins, including PURA/B, PCNA, TOP1, and PARP1, that all respond similarly to both EN^-^ and RT^-^ mutants compared to WT, exhibiting reduced associations with the mutant L1s; although, the RT^-^ mutant showed a larger effect size on the PURA/B proteins. These data suggest that cleavage of the host genomic DNA fosters associations between L1 and this group of proteins, but that interactions with PURA/B are further enhanced by L1 cDNA production. Other nuclear L1 proteins: HSPA8, HAX1, HSPA1A, TUBB, and TUBB4B were increased in both mutants. To better visualize the range of behaviors exhibited by our proteins of interest, and the population at large, we cross-referenced the relative enrichments of each protein detected in both experiments, shown in **Figure 3C**. We noted that the same striking trend mentioned above, two seemingly opposite behavioral classes of proteins could also be observed, globally among all proteins (**see Fig 3B, left side and right side, and Fig. 3C**). The crisscross pattern displayed in (and **Fig. S2**) indicates that two largely distinctive proteomes were purified with these two classes of mutants. Notably, the pattern observed appears to track with the relative behavior of ORF1p, which, along with other cytoplasmic L1 factors is elevated in RT^-^ mutants and reduced in EN^-^ mutants. We therefore speculate that the sum of observed interactomic changes include effects attributable directly to the catalytic mutations as well as indirect effects resulting from the response of ORF1p to the mutations.

**Figure 3.**
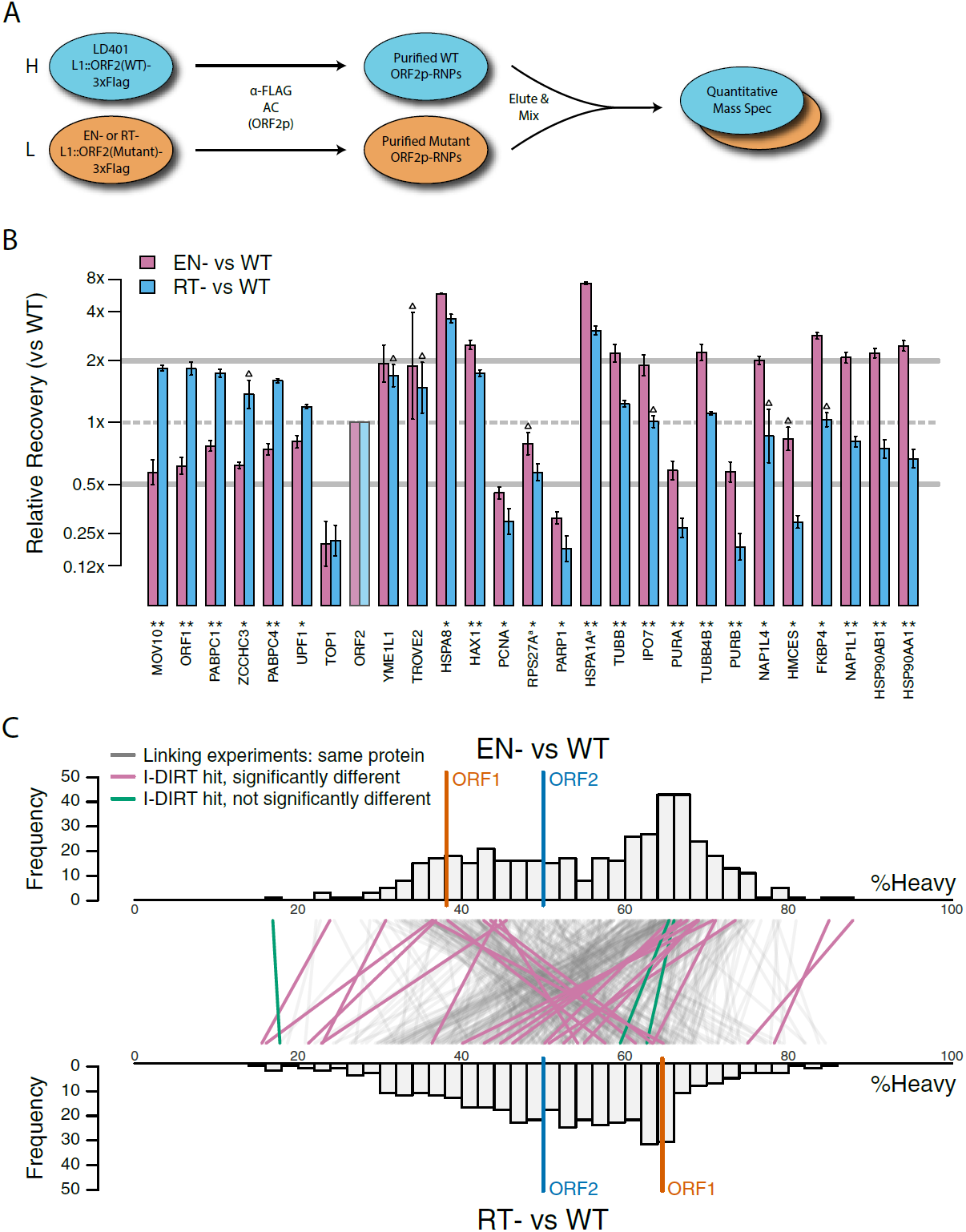
*Catalytic inactivation of ORF2p alters the L1 interactome*: L1s were affinity captured from cells expressing enzymatically active ORF2p-3xFLAG sequences (pLD401, WT), a catalytically inactivated endonuclease point mutant (pLD567; H230A, EN^---^), and a catalytically inactivated reverse transcriptase point mutant (pLD624; D702Y, RT). (**A**) WT L1s were captured from heavy-labeled cells, EN and RT L1s were captured from light-labeled cells. WT and either EN or RT fractions were mixed after purification, in triplicate, and the relative abundance of each co-purified protein in the mixture was determined by quantitative MS. (**B**) I-DIRT significant proteins displayed were detected in at least two replicates. All values were normalized to ORF2p. Data are represented as mean ± SD. Triangles (▵) mark proteins whose levels of co-purification did not exhibit statistically significant differences in the mutant compared to the WT. A single or double asterisk denotes a statistically significant difference between the relative abundances of the indicated protein in EN^-^ and RT^-^ mutants: p-values of between 0.05 -0.01 (*) and below 0.01 (**), respectively. Gray horizontal bars on the plot mark the 2x (upper) and 0.5x (lower) effect levels. (**C**) The double histogram plot displays the distributions of all proteins identified in at least two replicates, in common between both EN/WT (TOP) and RT/WT (LOWER) affinity capture experiments. The x-axis indicates the relative recovery of each copurifying protein and the y-axis indicates the number of proteins at that value (binned in 2 unit increments). The data are normalized to ORF2p. The relative positions of ORF2p and ORF1p are marked by colored bars. Differently colored lines illustrate the relative change in positions of the proteins within the two distributions (as indicated). Colored lines denote I-DIRT significance, with magenta lines indicating a statistically significant shift in position (p ≤0.05) within the two distributions and green lines indicating that statistical significance was not reached (entities labeled in **Fig. S2**). A cluster of magenta lines can be seen to track with ORF1p (red line, upper and lower histogram), and another cluster can be seen to behave oppositely, creating a crisscross pattern in the center of the diagram. A similar crisscross pattern is exhibited by many gray lines.

### 4.5 Dynamics of L1 RNPs *in vitro*

We next decided to measure the *in vitro* dynamics of proteins co-purifying with affinity captured L1s, reasoning that proteins with comparable profiles are likely candidates to be physically linked to one another or otherwise co-dependent for maintaining stable interactions with L1s. To achieve this, we first purified heavy-labeled, affinity-tagged L1s and subsequently incubated them, while immobilized on the medium, with light-labeled, otherwise identically prepared cell extracts from cells expressing untagged L1s (Luo *et al*, 2016). In this scenario, heavy-labeled proteins present at the zero time point are effectively "infinitely diluted" with light-labeled cell extract. The exchange of proteins, characterized by heavy-labeled proteins decaying from the immobilized L1s and being replaced by light-labeled proteins supplied by the cell extract, was monitored by quantitative MS. These experiments were conducted using constructs based on the naturally occurring L1RP sequence (Dai *et al*, 2014; Taylor *et al*, 2013; Kimberland *et al*, 1999). **Figure 4** illustrates the approach and displays the findings of our assay. We observed three distinctive clusters of behaviors (**Fig. 4B, C**). Notably, ORF1p, ZCCHC3, and the cytoplasmic poly(A) binding proteins clustered together, forming a relatively stable core complex. Exhibiting an intermediate level of relative *in vitro* dynamics, UPF1 and MOV10 clustered with TUBB, TUBB4B, and HSP90AA1. A third, and least stable, cluster consisted of only nuclear L1 interactors.

**Figure 4.**
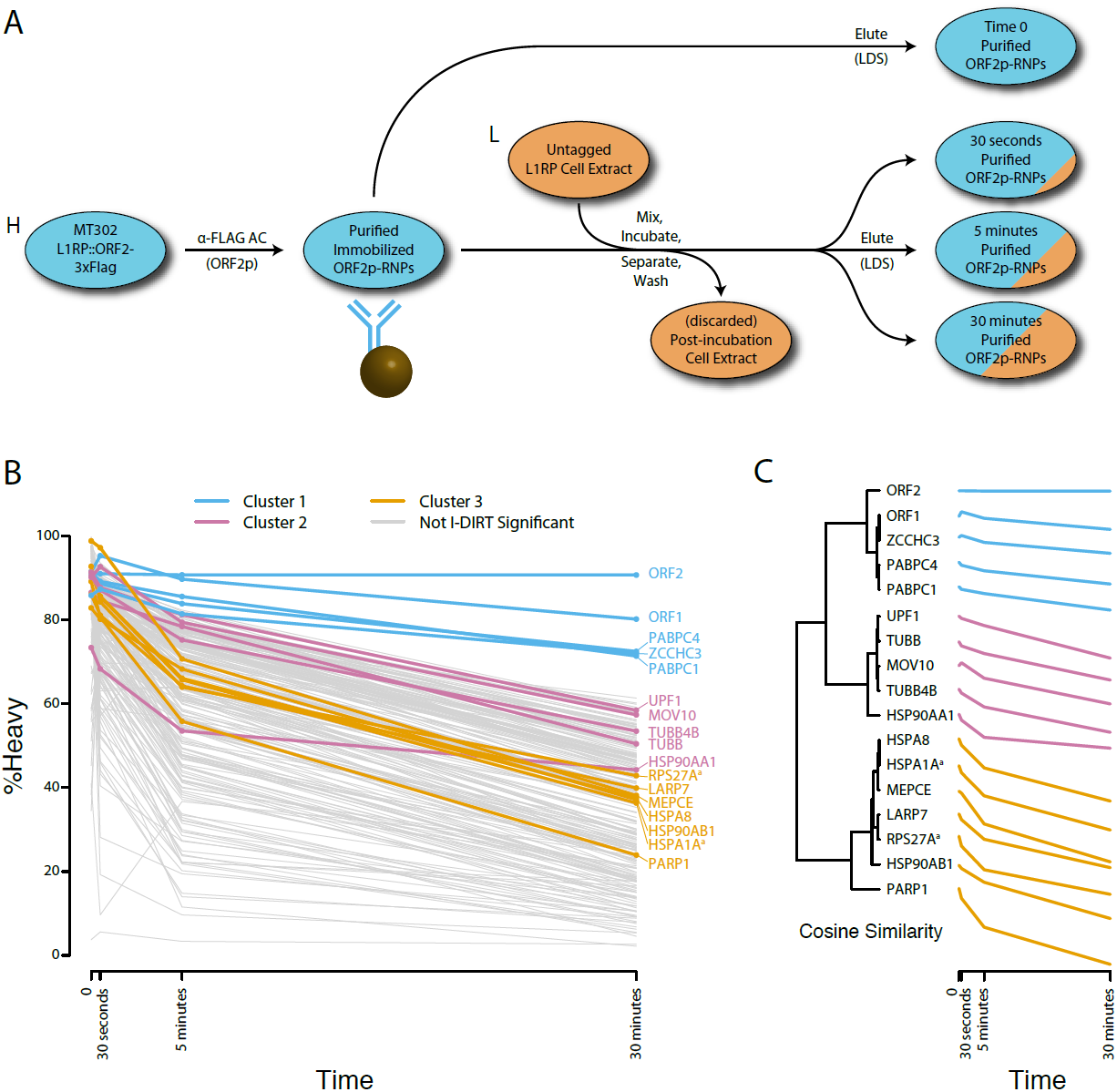
*Monitoring coordinated dissociation and exchange exhibited by L1 interactors in vitro*: L1s were affinity captured from heavy-labeled cells expressing ORF2p-3xFLAG in the context of the naturally occurring L1RP gene (pMT302); the stabilities of the protein constituents of the purified heavy-labeled L1 population were monitored *in vitro* by competitive exchange with light-labeled cell extracts containing untagged L1s (pMT298) (Taylor *et al*, 2013). (**A**) 3xFLAG-tagged L1s were captured from heavy-labeled cells and then, while immobilized on the affinity medium, were treated with an otherwise identically prepared, light-labeled, untagged-L1-expressing cell extract. Untreated complexes were compared to complexes incubated for 30 sec, 5 min, and 30 min, to determine the relative levels of exchange of *in vivo* assembled heavy-labeled interactors with *in vitro* exchanged light-labeled interactors using quantitative MS. (**B**) The results were plotted to compare the percentage of heavy-labeled protein versus time. I-DIRT significant proteins from **Table 1** are highlighted if present. Three clusters were observed (as indicated). (**C**) The cosine distance between the observed I-DIRT significant proteins was plotted along with time.

### 4.6 Multidataset Integration

Having observed coordinated and distinctive behaviors exhibited by groups of L1 interacting proteins across several distinctive biochemical assays, we then integrated the data and calculated the behavioral similarity of the I-DIRT-significant interactors, producing a dendrogram; **Figure 5** displays their relative similarities. A cluster containing the putative cytoplasmic L1 components (MOV10, UPF1, ZCCHC3, PABC1/4, ORF1p) was observed, as was a cluster containing PURA/B, PCNA, TOP1, PARP1, aligning with our assessments of the separated datasets (**Figs. 1, 3, 4**). In addition to these, we also observed three distinctive clusters derived from the nuclear L1 interactome. We believe that this is likely to reflect the presence of a collection of distinctive macromolecules.

**Figure 5.**
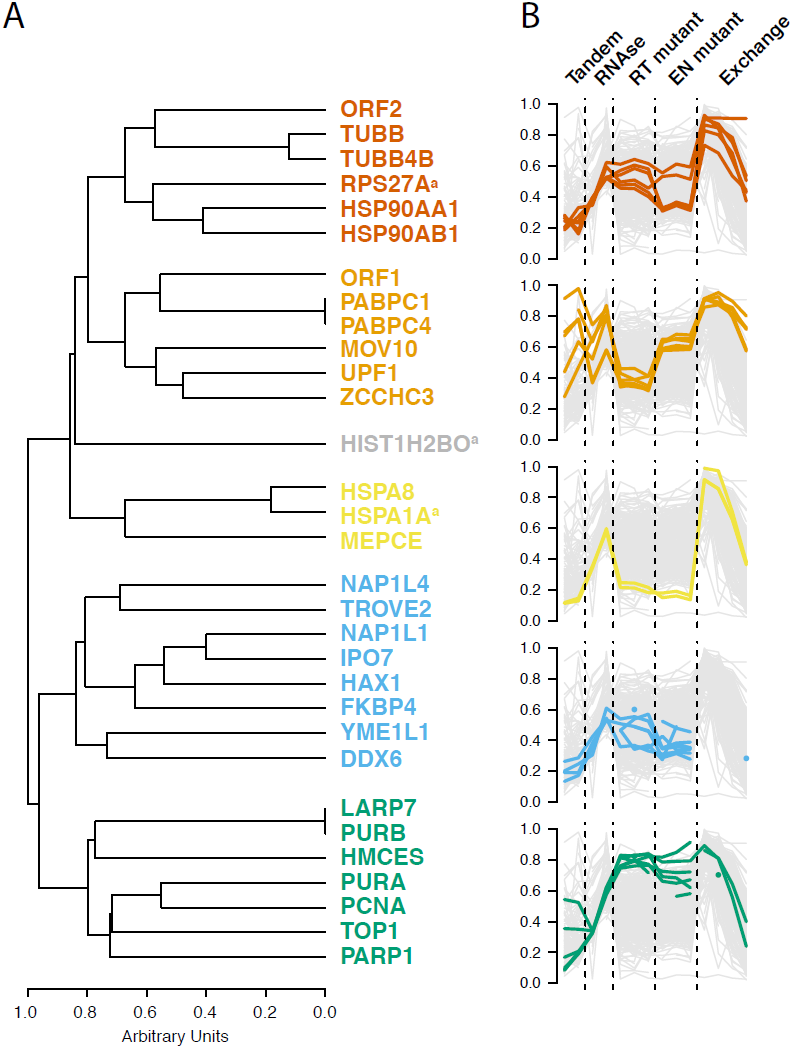
*Interactomic data integration*:(**A**) All MS-based affinity proteomic experiments presented were combined and analyzed for similarities across all I-DIRT significant proteins, producing five groupings. Distance are presented on a one-unit arbitrary scale. (**B**) The traces of each protein in each cluster, across all experiments, are displayed. The y-axis indicates the raw relative-enrichment value and the x-axis indicates the categories of each experiment-type. Each category is as wide as the number of replicates or time-point samples collected.

**Figure 6.**
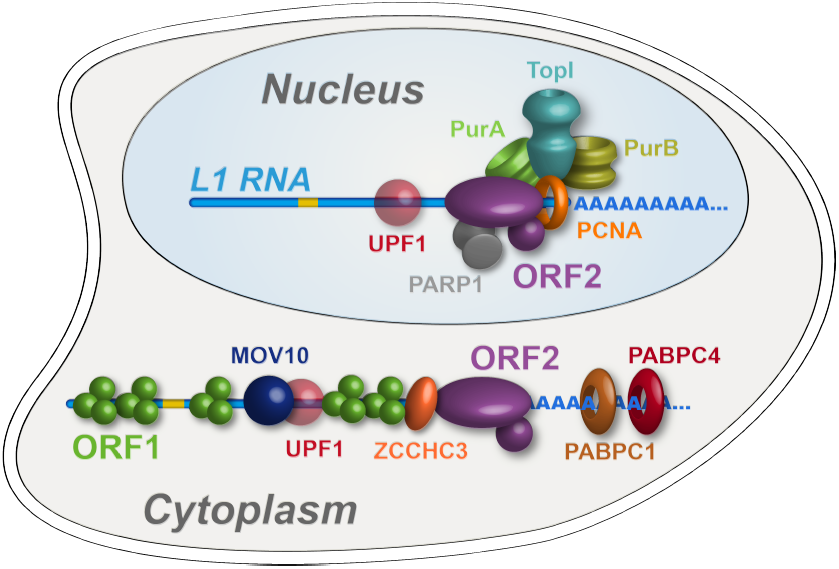
*Refined interatomic model*: Our results support the existence of distinct cytoplasmic and nuclear L1 interactomes. Affinity capture of L1 via 3xFLAG-tagged ORF2p from whole cell extracts results in a composite purification consisting of several macromolecular (sub)complexes. Among these, we propose a canonical cytoplasmic L1 RNP (**depicted**) and one or more nuclear macromolecules. UPF1 exhibited equivocal behavior within our fractionations and also co-purified with chromatin associated ORF2p, suggesting it participates in both cytoplasmic and nuclear L1 interactomes. Within the nuclear L1 interactome, our data support the existence of a physically linked entity consisting of PCNA, PURA/B, TOP1, and PARP1 (**depicted**).

## 5. Discussion

In this study we have characterized biochemical, interatomic, enzymatic, and cellular localization properties of ectopically expressed L1s. Through the assays explored, we observed discrete and coordinated behaviors, permitting us to refine our model of L1 intermediates, **diagrammed in Figure 7**. We propose a cytoplasmic L1, composed of ORF1/2p, L1RNA, PABPC1/4, MOV10, UPF1, and ZCCHC3, that constitutes an abundant, canonical RNP intermediate often referred to in the literature. We also propose a second more complex population, lacking ORF1p, that constitutes a nuclear L1 intermediate or, more likely, collection of intermediates. The nuclear population is enriched for factors linked to DNA replication and repair, including PURA, PURB, PCNA, TOP1, and PARP1; we propose that these proteins, along with ORF2p, form part of a direct intermediate of TPRT, although these components may not all act in synergy. Our proposals are broadly supported by the findings of **Mita et al. (co-submission)**, which also present data to support the hypothesis that PCNA-associated ORF2p is not appreciably associated with ORF1p, and also identified TOP1 and PARP1 in complex with ORF2p/PCNA.

### 5.1 Cytoplasmic L1s

ORF1p, MOV10, UPF1, and ZCCHC3 are released from L1 RNPs by treatment with RNases (**Fig. 1**), indicating the importance of the L1 RNA in the maintenance of these interactions. In this context, the L1 ORF and poly(A) binding proteins support L1 proliferation (Kulpa & Moran, 2006; Dai *et al*, 2012; Wei *et al*, 2001), whereas ZCCHC3 (**Table S2**) and MOV10 (Goodier *et al*, 2012; Arjan-Odedra *et al*, 2012) function in repressive capacities. Although UPF1 might also be expected to operate in a repressive capacity through its role in nonsense mediated decay (NMD), we previously demonstrated that UPF1’s role does not apparently resemble that of canonical NMD and it acts as an enhancer of retrotransposition despite negatively affecting L1 RNA and protein levels, supporting the possibility of repressive activity in the cytoplasm and proliferative activity in the nucleus (Taylor *et al*, 2013). Notably, MOV10 has been implicated in the recruitment of UPF1 to mRNA targets through protein-protein interactions (Gregersen *et al*, 2014). However, we observed that MOV10 exhibited a greater degree of RNase-sensitivity than UPF1, indicating that, if MOV10 directly modulates the UPF1 interactions with L1, a sub-fraction of UPF1 exhibits a distinct behavior (UPF1 is ~62% as sensitive to RNase treatment as MOV10, **Fig. 1C**). Bimodal UPF1 behavior can also be seen in split-tandem capture experiments, and UPF1 was recovered with L1s affinity purified from fractionated chromatin (**further discussed below**), and only about half of the UPF1 exhibits ORF1p-like partitioning with the canonical L1 RNP (**Fig. 1F**). Presumably, the RNase-sensitive fraction, released in concert with MOV10, is the same fraction observed in cytoplasmic L1s obtained by split-tandem purification. In contrast, PABPC1 and C4 exhibit strong ORF1p-like partitioning (comparable to MOV10), but appear wholly insensitive to RNase treatment. This is most likely due to the fact that neither RNase A nor T1 cleave RNA at adenosine residues (Volkin & Cohn, 1953; Yoshida, 2001); hence poly(A) binding proteins may not be ready targets for release from direct RNA binding by the assay implemented here (or generally, using these ribonucleases). Failure to release of ORF2p into the supernatant upon RNase treatment is expected due to its immobilization upon the affinity medium (Dai *et al*, 2014). However, we note that ORF2p binding to the L1 RNA has also been proposed to occur at the poly(A) tail (Doucet *et al*, 2015), raising the related possibility of close physical association on the L1 RNA between ORF2p and PABPC1/4 in cytoplasmic L1 RNPs. ORF1p, PABPC1/4, MOV10, ZCCHC3, and UPF1, all behaved comparably in response to EN-and RT-catalytic mutations, decreasing together in EN-mutants, and increasing together in RT-mutants (**Fig. 3B**). Moreover, when the exchange of proteins within L1 RNPs was monitored directly, PABPC1/4 and ZCCHC3 exhibited nearly identical stability, well above the background distribution; UPF1 and MOV10 also exhibited comparable kinetics to one another, falling into an intermediate stability cluster (**Fig. 4B, C**).

RNase-sensitivity was displayed by numerous proteins not previously identified as putative L1 interactors (**Table 1, Fig. 1** and (Taylor *et al*, 2013)). A known limitation of I-DIRT (and many SILAC-based analyses) is that it cannot discriminate non-specific interactors from specific but rapidly exchanging interactors (Wang & Huang, 2008; Luo *et al*, 2016; Smart *et al*, 2009). Our samples likely contain rapidly exchanging, physiologically relevant factors that were not revealed by I-DIRT under the experimental conditions used. With this in mind, we note members of the exon junction complex (EJC), RBM8A (Y14), EIF4A3 (DDX48), and MAGOH, are among our RNase-sensitive constituents, with all exhibiting a similar degree of RNase-sensitivity (**Fig. 1C**, labeled black dots). Crucially, these proteins are physically and functionally connected to UPF1 (reviewed in (Schweingruber *et al*, 2013)), and physically to MOV10 (Gregersen *et al*, 2014), both validated L1 interactors. We therefore hypothesize that EJCs may constitute bona fide L1 interactors missed in our original screen. This may seem unexpected because canonical L1 RNAs are thought not to be spliced, but this assumption has been challenged by one group (Belancio *et al*, 2006), and splicing-independent recruitment of EJCs has also been demonstrated (Budiman *et al*, 2009). Perhaps more compelling, EJC proteins exhibited a striking similarity in RNase-sensitivity to MOV10 (**Fig. 2B**). EIF4A3 has been suggested to form an RNA-independent interaction with MOV10 (Gregersen *et al*, 2014), and MOV10 is a known negative regulator of L1, making it attractive to speculate that these proteins were recruited and released in concert with MOV10 and/or UPF1.

Ectopically expressed canonical L1 RNPs have been shown to accumulate in cytoplasmic stress granules (Doucet *et al*, 2010; Goodier *et al*, 2010), and our observation of UPF1, MOV10, and MAGOH in the RNase-sensitive fraction is consistent with this characterization (Jain *et al*, 2016). However, the additional presence of EIF4A3 and RBM8A suggested that our RNPs may instead overlap with IGF2BP1 (IMP1) granules, reported to be distinct from stress granules (Jønson *et al*, 2007; Weidensdorfer *et al*, 2009). Consistent with this possibility, we observed IGF2BP1, YBX1, DHX9, and HNRNPU within the mixture of copurified proteins (**Table S1**). We did not, however, observe canonical stress granule markers G3BP1 or TIA1 (Goodier *et al*, 2007; Jain *et al*, 2016; Doucet *et al*, 2010). Surprisingly, siRNA knockdown of IGF2BP1 substantially reduced L1 retrotransposition; however, we note that the cytotoxicity associated with knocking-down EJC components may confound interpretation (**Table S2**). Given the result obtained, IGF2BP1 appears to support L1 proliferation. Consistent with an established function (Bley *et al*, 2015; Weidensdorfer *et al*, 2009), IGF2BP1 granules may sequester and stabilize L1 RNPs in the cytoplasm, creating a balance of L1 supply and demand that favors proliferation over degradation. Although human L1 does not contain a known IRES, it is known that *ORF2* is translated by a non-canonical mechanism (Alisch *et al*, 2006), and IGF2BP1 may promote this (Weinlich *et al*, 2009).

### 5.2 Nuclear L1s

The fraction eluted from the α-ORF1p medium contained the population of proteins physically linked to both ORF2p and ORF1p and greatly resembled the components released upon RNase treatment, hence these linkages primarily occur through the L1 RNA (or are greatly influenced by it). In contrast, the supernatant from the α-ORF1p affinity capture contained the proteins we speculate to be associated with ORF2p, but not ORF1p; moreover, fully intact RNA does not appear to be essential to the maintenance of these interactions. Notable within the latter group of interactors was the PURA/PURB/PCNA cluster, with TOP1 also in close proximity, ontologically grouping to the nuclear replication fork (GO:0043596). Separately, a few physical and functional connections have been shown for PURA/PURB (Knapp *et al*, 2006; Kelm *et al*, 1999; Mittler *et al*, 2009), PCNA/TOP1 (Takasaki *et al*, 2001), and PURA/PCNA (Qin *et al*, 2013). Notably, PURA, PURB, and PCNA have been independently linked to replication-factor-C / replication-factor-C-like clamp loaders (Kubota *et al*, 2013; Havugimana *et al*, 2012). Given that we also observe tight clustering of protein pairs known to be physically and functionally linked, e.g. PABPC1/4 (Jønson *et al*, 2007; Katzenellenbogen *et al*, 2007) and HSPA8/1A (Jønson *et al*, 2007; Nellist *et al*, 2005), and because we have established PCNA as a positive regulator of L1 retrotransposition (Taylor *et al*, 2013), we propose that the [PURA/B/PCNA/TOP1] group is a functional sub-complex of nuclear L1. In addition, although it does not cluster as closely to the [PURA/B/PCNA/TOP1] group, PARP1 is found within the putative nuclear L1 population and is functionally linked with PCNA, specifically stalled replication forks (Bryant *et al*, 2009; Min *et al*, 2013; Ying *et al*, 2016). Further tying them together, these proteins all also exhibited affinity capture yield decreases of severalfold in response to mutations that abrogated ORF2p EN or RT activity (**Fig. 3**). This is compelling because these ORF2p enzymatic activities are required in order for it to to manipulate DNA and traverse the steps of the L1 lifecycle that benefit from physical association with replication forks. One caveat to this interpretation is that, while knocking down PCNA reduced L1 retrotransposition (Taylor *et al*, 2013), no such effect was observed for TOP1 or PURA/B, which led instead to mild increases in L1 activity (**Table S2**). These proteins may be physically assembled within a common intermediate, but functionally antagonistic. HSP90 proteins were also observed in this fraction, and are also linked with stalled replication forks (Arlander *et al*, 2003; Ha *et al*, 2011), but exhibited a distinctive response to catalytic mutants, accumulating in EN-mutants while exhibiting a modest decrease in RT^-^ mutants. The recruitment of the ORF2p/PCNA complex to stalled replication forks has been also independently proposed by **Mita et al. (co-submission)**.

As mentioned above, we previously speculated that an RNase-insensitive fraction of L1-associated UPF1 may support retrotransposition in conjunction with PCNA in the nucleus ((Azzalin & Lingner, 2006; Taylor *et al*, 2013) and **Mita et al. [co-submission]**). In contrast to other PCNA-linked proteins, catalytic inactivation of ORF2p did not robustly affect the relative levels of co-purified UPF1, and UPF1 behaved in a distinct manner during tandem purification. The equivocal behavior of UPF1 in several assays (**Figs. 1, 3, and 4**) supports UPF1’s association with both the putative cytoplasmic and nuclear L1 populations, the latter being additionally supported by the association of UPF1 with ORF2p-3xFLAG purified from chromatin (**Table S3**). NAP1L4, NAP1L1, FKBP4, HSP90AA1, and HSP90AB1 (Baltz *et al*, 2012; Castello *et al*, 2012; Simon *et al*, 1994; Rodriguez *et al*, 1997; Peattie *et al*, 1992) are associated with RNA binding, involved in protein folding and unfolding, and function as nucleosome chaperones. An interesting possibility is that they have a nucleosome remodeling activity that may be required to allow reverse transcription to begin elongating efficiently, or for assembly of nucleosomes on newly synthesized DNA.

### 5.3 Future Studies

An obvious need is the continued validation of putative interactors by *in vivo* assays. RNAi coupled with L1 insertion measurements by GFP fluorescence (Ostertag *et al*, 2000) provides a powerful method to detect effects on L1 exerted by host factors. However, this approach can sometimes be limited by cell viability problems associated with important genes; it is therefore critical to control for this (**Table S2**). Along these lines, a high throughput approach has been developed for genome-wide screening of L1-host protein interactions (**P. Mita and J. Boeke, manuscript in preparation**). Throughout this and our prior study (Taylor *et al*, 2013) we have used comparable *in vitro* conditions for the capture and analysis of L1 interactomes. However, we are aware that this practice has enforced a single biochemical “keyhole” through which we have viewed L1-host protein associations. It is important to expand the condition space in which we practice L1 interactome capture and analysis in order to expand our vantage point on the breadth of L1-related macromolecules (Hakhverdyan *et al*, 2015). In concert with this, we must develop sophisticated, automated, reliable, low-noise methods to integrate biochemical, proteomic, genomic, and ontological data; the first stages of which we have attempted in the present study. Although we have used I-DIRT to increase our chances of identifying bona fide interactors (Tackett *et al*, 2005; Taylor *et al*, 2013), it is clear, and generally understood, that some proteins not making the significance cut-off will nevertheless prove to be critical to L1 activity (Byrum *et al*, 2011; Luo *et al*, 2016; Joshi *et al*, 2013), such as demonstrated by our unexpected findings with IGF2BP1 (**Table S2**). Through further development, including reliable integration with diverse, publicly available interactome studies, we hope to enable the detection of extremely subtle physical and functional distinctions between (sub)complexes and their components, considerably enhancing reliable exploration and hypothesis formation. Furthermore, it is striking that no structures of assembled L1s yet exist; these are missing data that are likely to provide a profound advance for the mechanistic understanding of L1 molecular physiology. However, we believe that with the methods presented here, endogenously assembled ORF1p/ORF2p/*L1-*RNA-containing cytoplasmic L1 RNPs can be prepared at sufficiently high purity and yield **Fig. 1F**) to enable electron microscopy studies. Importantly, we have shown that our affinity purified fractions are enzymatically active for reverse transcription of the L1 RNA (Taylor *et al*, 2013) and endonucleolytic cleavage of DNA (**J. LaCava, unpublished observation**), providing some hope that cryo-electron microscopy could be used to survey the dynamic structural conformations of L1s formed during its various lifecycle stages (Takizawa *et al*, 2017).

## 6. Methods

The preparation of L1 RNPs was carried out essentially as previously described (Taylor *et al*, 2013, 2016), with modifications described here. Briefly, HEK-293T_LD_ cells (Dai *et al*, 2012) transfected with L1 expression vectors were cultured as previously described or using a modified suspension-growth SILAC strategy described below. L1 expression was induced with with 1μg / ml doxycycline for 24 hours, and the cells were harvested and extruded into liquid nitrogen. In all cases the cells were then cryogenically milled (LaCava *et al*, 2016) and used in affinity capture experiments and downstream assays.

### 6.1 Modified SILAC Strategy

Freestyle-293 medium lacking Arginine and Lysine was custom-ordered from Life Technologies, and heavy or light amino acids plus proline were added at the same concentrations previously described (Taylor *et al*, 2013), without antibiotics. Suspension-adapted HEK-293T_LD_ were spun down, transferred to SILAC medium and grown for >7 cell divisions in heavy or light medium. On day 0, four (4) 1L square glass bottles each containing 200 ml of SILAC suspension culture at ~2.5×10^6^ cells/ml were transfected using 1 μg/ml DNA and 3 μg/ml polyethyleneimine "Max" 40 kDa (Polysciences, Warrington, PA, #24765). A common transfection mixture was made by pre-mixing 800 μg DNA and 2.4mL of 1 mg/ml PEI-Max in 40 ml Hybridoma SFM medium (Life Technologies, Grand Island, NY, #12045-076) and incubating for 20 min at room temperature (RT); 10 ml of the mixture was added to each bottle. On day 1, cells (200 ml) were split 1:2.5 (final two bottles each containing 250 mL) without changing the medium. On day 3, the cells were induced with 1 μg/ml doxycycline, and on day 4 the cells were harvested and extruded into liquid nitrogen. Aliquots were tested by western blot and the per-cell expression of both ORFs was indistinguishable from puromycin-selected material described previously (**Fig. S1**); transfection efficiency was assessed at >95% by indirect immunofluorescence of expressed ORF proteins. The median lysine and arginine heavy isotope incorporation levels for cell lines presented in this study were > 90%, determined as previously described (Taylor *et al*, 2013).

### 6.2 RNase-Sensitivity Affinity Capture

Four sets of 200 mg of light (L) and heavy (H) pLD401 transfected cell powders, respectively, were extracted 1:4 (w:v) with 20 mM HEPES-Na pH 7.4, 500 mM NaCl, 1% (v/v) Triton X-100 (extraction solution), supplemented with 1x protease inhibitors (Roche, Indianapolis, IN, #11836170001). After centrifugal clarification, all of the L and H supernatants were pooled, respectively, and then split, resulting in two sets of cleared L and H extracts equivalent to duplicate 400 mg samples from each SILAC cell powder. These four samples were each subjected to affinity capture upon 20 μl α-FLAG magnetic medium. After binding and washing, one set of L and H samples were treated with a control solution consisting of 2 μl of 2 mg/ml BSA (Thermo Fisher Scientific, Waltham, MA, #23209) and 50 μl extraction solution, v:v (Ctrl); the other set of L and H samples was treated with a solution of 2 μl 2 mg/ml RNase A / 5000 u/ml RNase T1 (Thermo Fisher Scientific #EN0551) and 50 μl extraction solution, v:v (RNase). Samples were then incubated 30 min at RT with agitation, the supernatant was removed, and the medium was washed three times with 1 ml of extraction solution. The retained captured material was eluted from the medium by incubation with 40 μl 1.1x LDS sample loading buffer (Life Technologies #NP0007). To enable quantitative comparisons of fractions, the samples were combined, respectively, as follows: 30ul each of the ^LL^RNase with ^HH^Ctrl, and 30ul each of the Ctrl with RNase. These samples were reduced, alkylated and run until the dye front progressed ~6 mm on a 4-12% Bis-Tris NuPAGE gel (Life Technologies, as per manufacturers instructions). The gels were subsequently subjected to colloidal Coomassie blue staining (Candiano *et al*, 2004) and the sample regions ("gel-plugs") excised and processed for MS analyses, as described below.

### 6.3 Split-Tandem Affinity Capture

400 mg of light (L) and heavy (H) pLD401 transfected cell powders, respectively, were extracted and clarified as above. These extracts were subjected to affinity capture on 20 μl α-FLAG magnetic medium, 30 min at 4°C, followed by native elution with 50 μl 1 mg/ml 3xFLAG peptide (15 min, RT). 45 μl of the elution were subjected to subsequent affinity capture upon 20 μl α-ORF1 magnetic medium, resulting in a 45 μl supernatant (Sup) fraction depleted of ORF1p. Finally, the material was eluted (Elu) from the α-ORF1p medium in 45 μl 2.2x LDS sample loading buffer by heating at 70°C for 5 min with agitation. To enable quantitative comparisons of fractions the samples were combined, respectively, as follows: 28 μl each of the ^L^Sup with ^H^Elu, and 28 μl each of the ^L^Elu with^H^Sup. These samples were then prepared as gel-plugs (as above) and processed for MS analyses, as described below.

### 6.4 Mass Spectrometry Sample Preparation and Data Acquisition

Gel plugs were excised, cut into 1 mm cubes, de-stained, and digested overnight with enough 3.1 ng/μl trypsin (Promega, Madison, WI, #V5280) in 25 mM ammonium bicarbonate to cover the pieces. In RNase-sensitivity and split-tandem SILAC analyses based on pLD401, as well as *in vitro* protein exchange experiments based on pMT302 and pMT289, an equal volume of 2.5 mg/ml POROS R2 20 μm beads (Life Technologies #1112906) in 5% v/v formic acid, 0.2% v/v TFA was added, and the mixture incubated on a shaker at 4°C for 24 hr. Digests were desalted on Stage Tips (Rappsilber *et al*, 2007), eluted, and concentrated by vacuum centrifuge to ~10 μl. ~3 μl were injected per LC-MS/MS analysis. RNase-sensitivity and split-tandem samples were loaded onto a PicoFrit column (New Objective, Woburn, MA) packed in-house with 6 cm of reverse-phase C18 material (YMC*Gel ODS-A, YMC, Allentown, PA). Peptides were gradient-eluted (Solvent A = 0.1 M acetic acid, Solvent B = 0.1 M acetic acid in 70% v/v acetonitrile, flow rate 200 nl/min) into an LTQ-Orbitrap-Velos or an LTQ-Orbitrap-XL mass spectrometer (Thermo Fisher Scientific) acquiring data-dependent CID fragmentation spectra. *In vitro* exchange samples were loaded onto an Easy-Spray column (ES800, Thermo Fisher Scientific) and gradient-eluted (Solvent A = 0.1% v/v formic acid in water, Solvent B = 0.1% v/v formic acid in acetonitrile, flow rate 300 nl/min) into an Q Exactive Plus mass spectrometer (Thermo Fisher Scientific) acquiring data-dependent HCD fragmentation spectra. In SILAC experiments comparing inactivated ORF2p catalytic mutants to WT (based on pLD401 [WT], pLD567 [EN-], and pLD624 [RT-]) peptides were extracted from the gel in two 1 hr incubations with 1.7% v/v formic acid, 67% v/v acetonitrile at room temperature with agitation. Digests were partially evaporated by vacuum centrifugation to remove acetonitrile, and the aqueous component was desalted on Stage Tips. Peptides were loaded onto an Easy-Spray column (ES800, Thermo Fisher Scientific) and gradient-eluted (Solvent A = 0.1% v/v formic acid in water, Solvent B = 0.1% v/v formic acid in acetonitrile, flow rate 300 nl/min) into an Orbitrap Fusion Tribrid mass spectrometer (Thermo Fisher Scientific) acquiring data-dependent fragmentation spectra (either CID spectra alone, or CID and HCD spectra).

### 6.5 Mass Spectrometry Data Analysis

Raw files were submitted to MaxQuant (Cox & Mann, 2008) version 1.5.2.8 for protein identification and isotopic ratio calculation. Searches were performed against human protein sequences (UP000005640, April 2016), custom L1 ORF1p and ORF2p protein sequences, common exogenous contaminants, and a decoy database of reversed protein sequences. Search parameters included fixed modification: carbamidomethyl (C); variable modification: Arg10, Lys8, methionine oxidation; razor and unique peptides used for protein quantitation; requantify: enabled. Contaminants, low-scoring proteins and proteins with one razor+unique peptides were filtered out from the MaxQuant output file "proteingroups.txt". The list of contaminants was uploaded from the MaxQuant web-site (http://www.coxdocs.org/; "contaminants"). Additionally, proteins with the "POTENTIAL CONTAMINANT" column value "+" were filtered out. Proteins with at least 2 razor+unique peptides were retained for the analysis. H/(H+L) and L/(H+L) values were derived from unnormalized "ratio H/L" values and were used for plotting label-swapped RNase-sensitivity and split-tandem data. Unnormalized "ratio H/L" values were used to calculate H/(H+L) in ORF2p catalytic mutant comparisons and in vitro exchange experiments. These values have been referred to as "affinities" within the **Supplementary Materials**. Normalization and clustering procedures applied to data presented in the figures (**Table S1**) are detailed below and also in the **Supplementary Methods**.

To plot RNase-sensitivity affinity capture results (**Fig. 1C**), these data were normalized such that proteins that did not change upon treatment with RNases are centered at the origin. The mean value and standard deviation were calculated using the distribution of distances from the origin. The distance threshold for p-value = 0.001 was calculated using the R programming language. A circle with radius equal to the threshold was plotted. Points with distances higher than the threshold were marked as black. To plot split-tandem affinity capture results (**Fig. 1F**), these data were normalized such that the ORF1p affinity was set to 1 and the distribution median was maintained. Probabilities associated with selected clusters were calculated based on the frequency distributions of 2-and 3-node clusters present in the data. To plot EN^-^ and RT^-^ mutant affinity capture results (**Fig. 3B**), the matrix of detected proteins for each experiment (EN^-^ and RT^-^) was filtered to retain only proteins detected in at least two replicate experiments. The difference between the affinity value of ORF2p and 0.5 value was calculated for each experiment. The affinities of each protein were shifted by the calculated difference. To determine the statistical significance of differentially co-purified proteins between EN^-^ or RT^-^ and WT, respectively, we used a 1-sample t-test and applied Benjamini-Hochberg p-value correction. To determine the statistical significance of differentially co-purified proteins between EN^-^ and RT^-^ we used an unpaired t-test and applied Benjamini-Hochberg p-value correction. To plot *in vitro* dynamics (**Fig. 4B, C**), only proteins which were identified at all time points were used. The cosine similarity method was used to calculate distances between proteins, and hierarchical clustering was used to visualize these distances. To integrate and plot the combined data (Fig. 5), we calculated Euclidean and cosine distances for each I-DIRT-significant protein pair present in each experiment. Euclidean distances were rescaled to the range (0, 0.9). Proteins not detected in any common experiments were assign a Euclidian distance of 1 after rescaling. The total distance between protein pairs was calculated as d = log((Rescaled Euclidean distance) * (cosine distance)). This distance was rescaled to the range (0, 1). Hierarchical clustering was used to visualize the calculated distances.

### 6.6 Gene Ontology (GO) Analysis

Genes corresponding to the proteins previously reported as significant by I-DIRT (Taylor *et al*, 2013) were tested for statistical overrepresentation using the default settings provided by http://www.panthnerdb.org (Mi *et al*, 2017, 2013), searches were conducted using GO complete molecular function, biological process, and cellular compartment: all results are compiled in **Table S4**.

### 6.7 ORF protein immunofluorescence analysis in HeLa cells

Tet-on HeLa M2 cells (Hampf & Gossen, 2007) (a gift from Gerald Schumann), were transfected and selected with 1 μg/ml puromycin for three days. Puromycin-resistant cells were plated on coverslips pre-coated for 1-2 hr with 10 μg/ml fibronectin in PBS (Life Technologies). 8-16 hr after plating, L1 was induced with [inline] doxycycline. 24 hr later, cells were fixed in 3% paraformaldehyde for 10 min. Fixative was then quenched using PBS containing 10 mM glycine and 0.2% w/v sodium azide (PBS/gly). The cells were permeabilized for 3 min in 0.5% Triton X-100 and washed twice with PBS/gly. Staining with primary and secondary antibodies was done for 20 min at room temperature by inverting coverslips onto Parafilm containing 45ml drops of PBS/gly supplemented with 1% BSA, mouse anti-FLAG M2 (Sigma, 1:500), rabbit anti-ORF1 JH73 (1:4000) (Taylor *et al*, 2013), Alexa Fluor 488 conjugated anti-mouse IgG (Life Technologies, 1:1000), and Alexa Fluor 568 conjugated anti-rabbit IgG (Life Technologies, 1:1000). DNA was stained prior to imaging with Hoechst 33285 (Life Technologies, 0.1 μg/ml). Epifluorescent images were collected using an Axioscop microscope (Zeiss, Jena, Germany) equipped for epifluorescence using an ORCA-03G CCD camera (Hamamatsu, Japan).

## 7. Author Contributions

Conceptualization, J.L., M.S.T., J.D.B.; Methodology, J.L., K.R.M., M.S.T., I.A.; Software, I.A.; Formal Analysis, I.A., D.A., D.F., D.I.; Investigation, J.L., K.R.M., M.S.T., P.M., H.J.,.M.A., A.W.; Resources, M.P.R., B.T.C., D.A., J.D.B., D.F., K.H.B.; Data Curation, K.R.M., I.A., J.L., D.A.; Writing -Original Draft, J.L., M.S.T.; Writing -Review & Editing, J.L., M.S.T., P.M., M.P.R., J.D.B., K.H.B.; Visualization, I.A., J.L., M.S.T., D.I.; Supervision, J.L., D.A.; Project Administration, J.L.; Funding Acquisition, M.P.R., B.T.C., J.D.B., D.A., K.H.B.

## Acknolwedgements

This work was supported in part by National Institutes of Health (NIH) grants P41GM109824 (to M.P.R.), P41GM103314 (to B.T.C.), and P50GM107632 (to J.D.B), and by the 5-100 Russian Academic Excellence Program. The mass spectrometric analysis of proteins co-purified with chromatin associated ORF2p (**Table S3**) was conducted within the NYU School of Medicine Proteomics Resource Lab, which is partially supported by the Laura and Isaac Perlmutter Cancer Center Support Grant, NIH P30CA16087, and NIH 1S10OD010582. This paper is subject to the NIH Public Access Policy. The authors declare no conflicts of interest.

